# Decrease in within-trial variability contributes to a decrease in across-trial variability of neural firing in the primate cortex during neural computations

**DOI:** 10.1101/635532

**Authors:** Naveen Sendhilnathan, Debaleena Basu, Aditya Murthy

## Abstract

The conventional approach to understanding neural responses underlying complex computations is to study across-trial averages of repeatedly performed computations from single neurons. When a brain region performs complex computations, such as processing stimulus related information or motor planning, it has been repeatedly shown through measures such as the Fano factor (FF) that neural variability across trials in the network decreases. However, in reality, multiple neurons contribute to a common computation on a single trial, rather than a single neuron contributing to a computation on multiple trials. Therefore, on individual trials the concept of FF loses significance. In this study, we extended previous work using measures of variability that are confined to a single trial and found that neurons perform a common computation, they briefly fire with increased regularity in spike timings, with similar inter-spike interval durations. We propose that this decrease in within-trial variability in neural spiking contributes to a decrease in across-trial variability in neural firing rates during network level computations. We confirmed our hypothesis by testing it on the activity of frontal eye field neurons recorded as two monkeys performed a memory-guided saccade task, and also on simulated spike trains. Furthermore, this phenomenon also has important behavioral correlates: the reaction time of the animal was faster when the within-trial variability was lower. We show that a decrease in within-trial variability is linked to a decrease in across-trial variability in neural response and indicates stationarity of neural network variability across time.

**New & Noteworthy:** During computations, neural variability across trials decreases. In reality, multiple neurons contribute to a common computation on a single trial, rather than a single neuron contributing to a computation on multiple trials. We found that when a network of neurons performs a common computation, they briefly fire with increased regularity in spike timings. We propose that this decrease in within-trial variability in neural spiking contributes to a decrease in the observed across-trial variability.

## Introduction

Neurons process information and perform computations through spikes. However, the pattern of neural activity is surprisingly variable, even across trials where the same stimulus is presented repeatedly, and with similar stimulus-response mappings ^1,2^ and the behavior tightly controlled. Given the inherent stochastic nature of neural activity, there is a long-standing debate as to whether the information in the spike content is encoded in the rate of spikes, or the precise timing of spikes ^3^. The conventional approach to studying neural activity is through trial averaging the spike frequency and considering that the firing rate bears the kernel of neural computation, disregarding the trial-to-trial spiking variability under similar experimental conditions, as random fluctuations or ‘noise’ in the system.

For a single neuron, spiking noise or variability can be characterized using different time-scales: ‘within-trial’ variability, reflecting the randomness in the temporal sequence of action potentials within single trials, and ‘across-trial’ variability, referring to the changes in spike counts over multiple trials. Previous studies have established that the across-trial variability in neural firing decreases during neural computations such as in response to the onset of a stimulus, suggesting the possibility that neural computations suppress the chaos in the system making it more ‘stable’ following an input ^4,5^. Decrease in neuronal variability has also been correlated with broad spatial tuning after stimulus onset ^6^, expected reward^7^, attention^8^, task familiarity ^9^, sensorimotor learning ^10^, motor preparation^11^, task engagement ^12^ and behavior ^4,7,13,14^. The average across-trial neuronal variability is considered a proxy for the state of the neural network that performs population level coding ^15,16^ and is correlated with local field potential oscillations ^17^. However, such fluctuations that are present only across several trials are unlikely to be measured or used by the brain towards any computation on a given single trial. This is because multiple neurons contribute to a common computation on a single trial, rather than a single neuron contributing to a computation on multiple trials. Therefore, while task-related modulations in the across-trial variability has been explored by previous studies, the question of how within-trial variability influences across-trial variability still remains to be answered^4^.

Consider the scenario presented in Fig 1. The input driven spiking activity for three trials, for this given neuron, could have Poisson like spiking variability at the scale of single trials (Fig 1a), or could be highly regular for a brief period (Fig 1b). Furthermore, there could be a third scenario where within any given trial, the spiking could be regular; however, this ‘magnitude of regularity’ can be different for different trials. Notwithstanding the difference in variability, the firing rates in all these conditions can be identical, with similar spike density functions. We propose that the across-trial variability decreases only in the second case and not in the former or the latter. In other words, we propose that during a neural computation, the neurons fire with increased regularity and this decrease in the within-trial variability in spiking contributes to a decrease in across-trial variability in firing rate.

**Figure 1.**
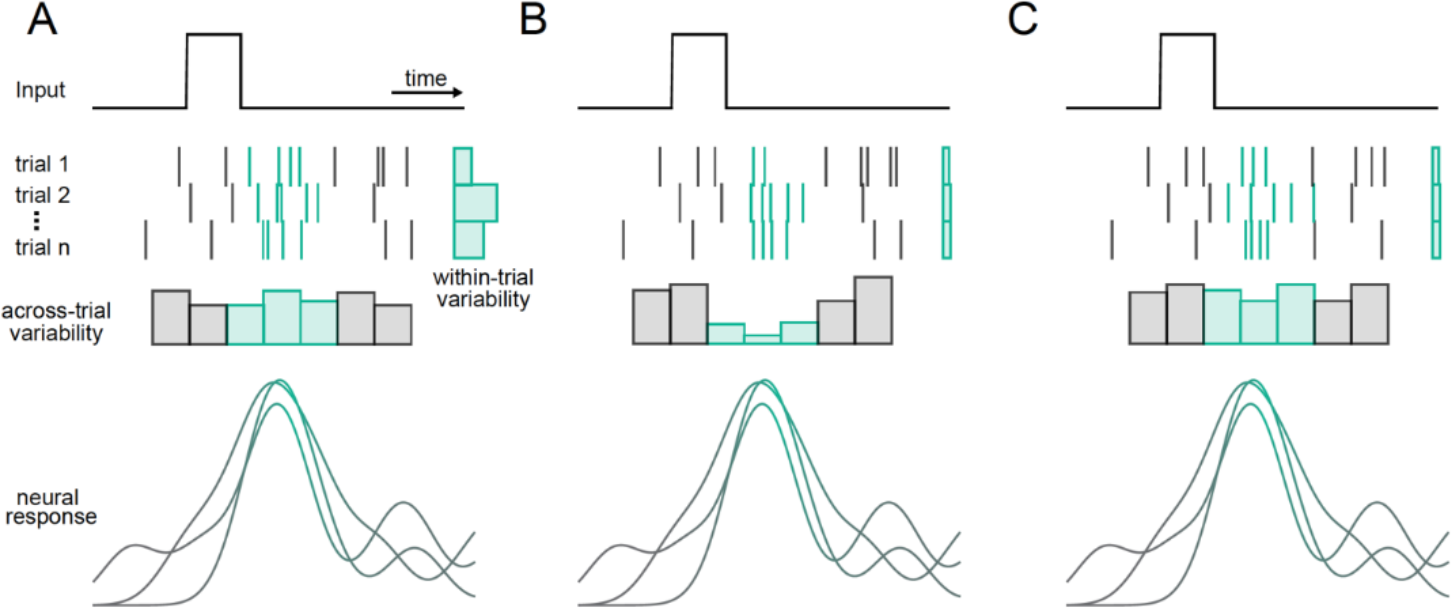
top panel: Schematic illustration of three possible scenarios where the spike density functions (bottom panel) in response to an input (top panel) are the same, but the inter-spike intervals are different (middle panel). **A**: The within-trial variability in firing is high during a small epoch (marked by green rasters), modelled as a Poisson firing model (gamma distribution shape parameter, k=1). **B**: The within-trial variability in firing is low during a small epoch (marked by green bars in the bottom), modelled as regular firing model (gamma distribution shape parameter, k=4). The extent of inter-trial regularity is similar across trials. **C:** The within-trial variability in firing is low during a small epoch (marked by green bars in the bottom) in individual trials, modelled as a within-trial regular firing model (gamma distribution shape parameter, k=4), but the extent of inter-trial regularity is different across trials.

We tested our hypothesis on a dataset of frontal eye field (FEF) neurons from two monkeys acquired as they performed a memory-guided saccade task ^18^. We then simulated neurons with different levels of within-trial variability and firing rates to confirm our hypothesis. Finally, following up on previous results that faster reaction times (RT) correlate with lesser across-trial variability, we reasoned that if within-trial variability does drive the variability across trials, it should correlate similarly with RT and that is what we observed. Thus, taken together, our results suggest that at the level of single trials, the neurons fire with increased regularity in spike timings while performing task-driven neural computation and that this decrease in within-trial variability can contribute to a decrease in across-trial variability when seen at the scale of multiple trials.

## Results

We performed all our analyses partly on a dataset of frontal eye field neurons that was published earlier ^18^ and partly on simulated neurons (see methods). We analyzed 70 FEF neurons that were recorded from two monkeys as they were performing a memory-guided saccade task where the monkeys had to remember precisely, one of the eight target locations where a target was flashed briefly, and then make a saccade ~1000ms later, to the remembered location to earn a liquid reward (see methods for details). The memory-guided saccade task requires successful encoding of the target stimulus, maintaining the location in memory during the delay period, and thereafter making a saccade to the remembered location, thus allowing for separation of visual, delay, and saccade related processes in the FEF and assessing how the two types of variability change in these three epochs.

### Across-trial variability decreased during a target presentation and movement preparation

We quantified the across-trial variability in neural firing using the Fano factor (FF), which is defined as the variance in the signal normalized by the mean. This provided an estimate of the ‘noise-to-signal’ ratio and captured the normalized amount of variability in the signal. The spiking noise is roughly assumed and modelled as a Poisson process. Since the variance of a Poisson distribution scales with its mean, the FF mathematically equals one for a Poisson process. Thus, a decrease in across-trial variability could be measured as a decrease in variance, compared to its mean. In our dataset, relative to the FF in the fixation epoch (1.145±0.0123), the FF in the target epoch (Fig 2B) and the saccade epoch (Fig 2D) decreased significantly in the receptive field (RF; target epoch: P = 1.3491e-08; ranksum test; saccade epoch: P = 1.0237e-18; ranksum test) and only slightly decreased in position 180 degrees opposite to the RF (aRF; target epoch: P=0.0265, ranksum test; saccade epoch: P = 2.0420e-05; ranksum test), resulting in a slight spatial modulation ^6^. This decrease in FF was absent for both RF (P = 0.9926; ranksum test) and aRF (P = 0.6050; ranksum test) conditions in the delay epoch (Fig 2C).

**Figure 2.**
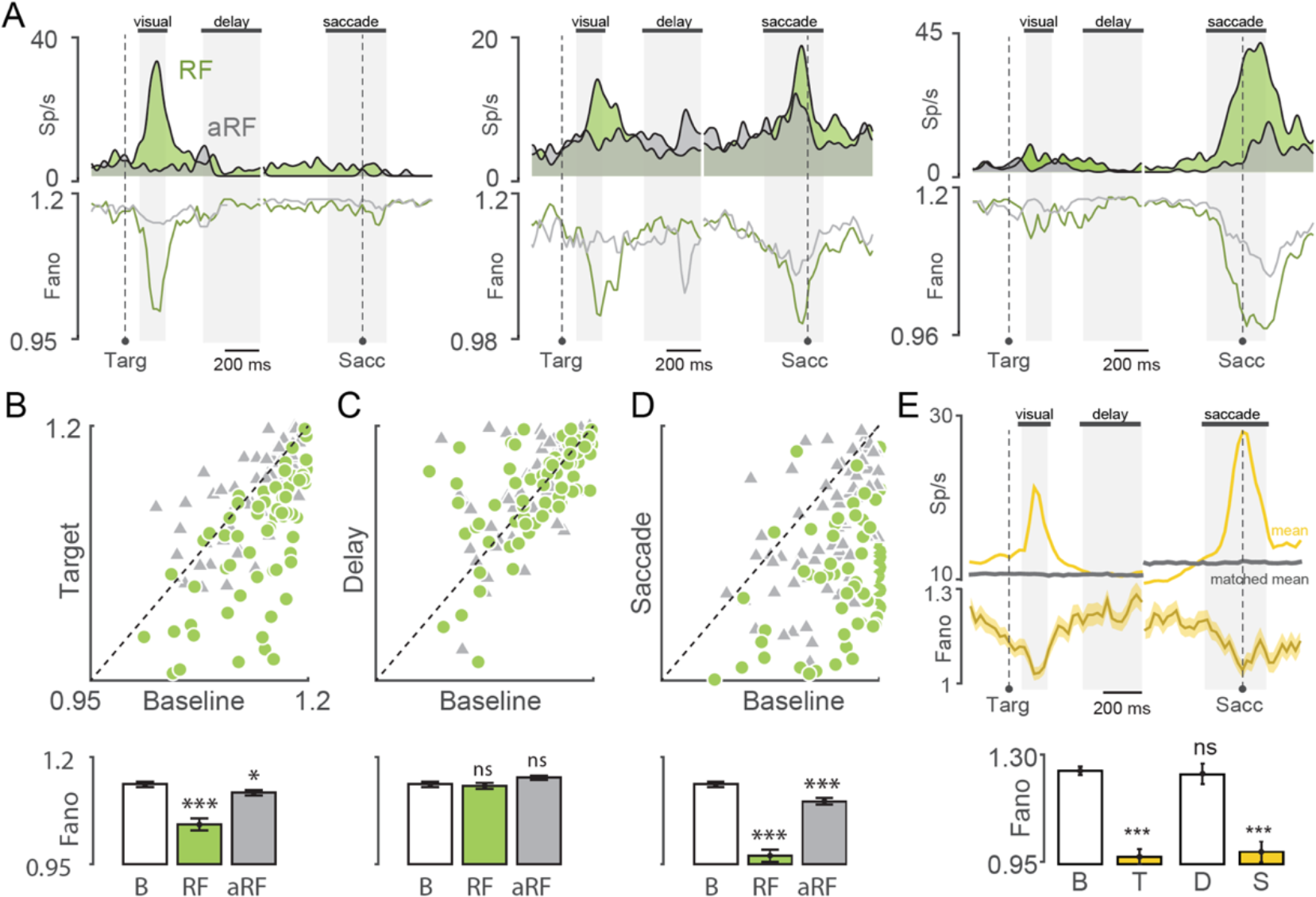
Across-trial variability decreased during a stimulus driven response and movement preparation: **A.** Top: Spike density function of a representative FEF visual (left), visuomotor (center) and motor (right) neuron during target epoch and saccade epoch for receptive field (green) and anti-receptive field (grey). Bottom: FF for the same neurons for receptive field (green) and anti-receptive field (grey). **B.** top panel: FF computed for RF (green circles) and aRF (grey triangles) positions for all neurons in the target epoch plotted against the fixation epoch. The broken diagonal line is the line of unity. Bottom panel: bar plot of mean FF in the fixation epoch (B) and target epoch in RF and aRF. *** means P<0.001; * means P<0.05, ranksum test. **C.** Same as B, but for delay epoch. n.s means not significant, ranksum test. **D.** Same as B, but for saccade epoch. *** means P<0.001; *** means P<0.001, ranksum test. **E.** top panel: Mean population neural response across all conditions during target epoch and saccade epoch (top yellow traces). Mean-matched neural responses (grey) and mean-matched FF (bottom yellow) traces computed using method described in 4. Bottom panel: bar plot of average mean-matched FF in the fixation epoch (B) and target epoch (T), delay (D) and saccade epoch (S). *** means P<0.001; n.s means not significant, ranksum test. All the P values reported were from appropriate statistical tests, tested against the corresponding fixation values.

However, during neural computations, since the firing rate usually increases, the FF tends to decrease trivially, by definition, providing an incorrect readout of the variability in neural firing. To circumvent this problem, we used the technique developed by Churchland et.al., where we computed FF on the mean-matched firing rates ^4^. We computed mean-matched FF in the fixation epoch, target epoch, delay epoch and the saccade epoch for all three types of FEF neurons together (Fig 2E) and observed the same results. Thus, confirming prior results, we report that the across-trial variability decreased both during target epoch and the saccade epoch in FEF neurons ^13,19^. Henceforth, we will collectively refer to both target onset, and movement preparation-execution as neural events.

### Within-trial variability decreased during a target presentation and movement preparation

Although a measure of the across-trial variability provides a platform to look at the network dynamics during neural computation on the scale of multiple trials, it loses its significance on a single trial basis. Multiple neurons contribute to a common computation on a single trial, rather than a single neuron contributing to a computation on multiple trials. Therefore, we looked at the pattern of neural activity during neural events within single trials, that could explain the reduction of variability across trials.

The variability on a single trial level comes from the variability of spike timings. Standard methods such as coefficient of variation of interspike intervals (ISI; standard deviation normalized by mean) are subject to variation in firing rate. Therefore, we measured the variability in spike timings by characterizing the variance of ISI: small variability in ISI implies regular spiking and high variability implies irregular spiking (Fig 3A). We used the CV2 index, which is defined as the difference in consecutive ISIs normalized by their mean to measure this variability in ISI ^20^. This provides a non-parametric measure of the regularity in spiking within a trial. We found that, the CV2 relative to the fixation epoch (0.976±0.032), decreased significantly in the RF in the target epoch (P = 0.0066; ranksum test, Fig 3B) and the saccade epoch (P = 0.0056; ranksum test, Fig 3D) and only slightly decreased in the aRF (Target epoch: P = 0.0187, ranksum test Fig 3B; Saccade epoch: P = 0.04513 ranksum test, Fig 3D), while there was no modulation in the delay epoch (RF: P = 0.2461, ranksum test; aRF: P = 0.6765, ranksum test; Fig 3C), echoing the patterns observed with FF. Clearly, a decrease in CV2 indicates increased regularity of spiking, thus a decreased within-trial variability in neural firing.

**Figure 3.**
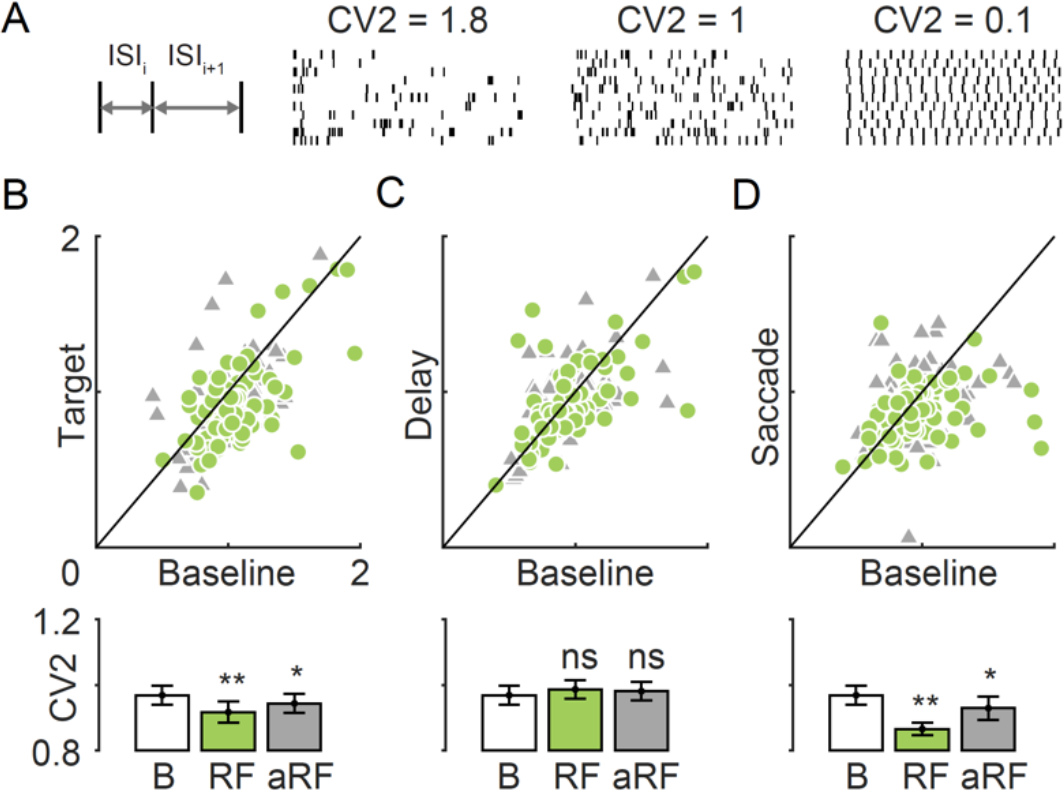
Within-trial variability in firing decreased in the target and saccade epochs but not in the delay epoch. **A.** Schematic illustration of CV2 calculation. Left: CV2 estimates the variation between the two ISIs of three consecutive spikes. Right: three simulated rasters with different CV2 values as indicated in top showing their spiking properties. A lower CV2 indicates higher level of spike regularity and low within-trial variability in spiking. **B.** Top: CV2 in the target epoch for RF (green circle) and aRF (grey triangle) sides during fixation versus target epoch. Diagonal line is the line of unity. Each pair of markers (circle and triangle) is a neuron. Bottom: mean FF in the fixation (white) and the target epoch for RF (green) and aRF (grey) sides. ** means P<0.01, and * means P<0.05, ranksum test. **C.** Same as B, but for delay epoch. n.s means not significant, ranksum test. **D.** Same as B, but for saccade epoch. ** means P<0.01, ttest. * means P<0.05, ranksum test.

Although CV2 provides an estimate of spiking variability in short intervals, it is not entirely reliable when considering long range slow changes in firing rate. Therefore, to verify that our results were not biased, we used another method, that is independent of firing rate fluctuations. First, we generated ISI histograms for each of the four epochs (fixation, target, delay and saccade) for each neuron and then fit a gamma function ^21–23^ to the histograms (see methods). Gamma distributions are defined by a shape parameter (k) providing a measure of ISI regularity and a scale parameter (θ), approximately providing an estimate of firing statistics. For instance, if k=1, the gamma distribution reduces to a Poisson model. If k is different from 1, it indicates a deviation from Poisson firing with k<1 signifying ISIs being too short and too long, and k>1 signifying increased regularity in firing. Hence, the shape parameter of the gamma fit is a convenient measure of spiking regularity and within-trial variability in neural firing. The ISI histograms, across different conditions, were well described as gamma distributions. Confirming our earlier observation, we found that the k values for the distribution of ISIs, compared to the fixation epoch (k = 1.53 ± 0.10), were significantly greater than one during target (k = 2.33 ± 0.19; P = 6.2953e-06; ranksum test) and movement (k = 2.29 ± 0.15; P = 2.1799e-07; ranksum test) epochs but not during the fixation or delay (k = 1.72 ± 0.20; P = 0.5958; ranksum test) epochs (Fig 4 A-C).

**Figure 4.**
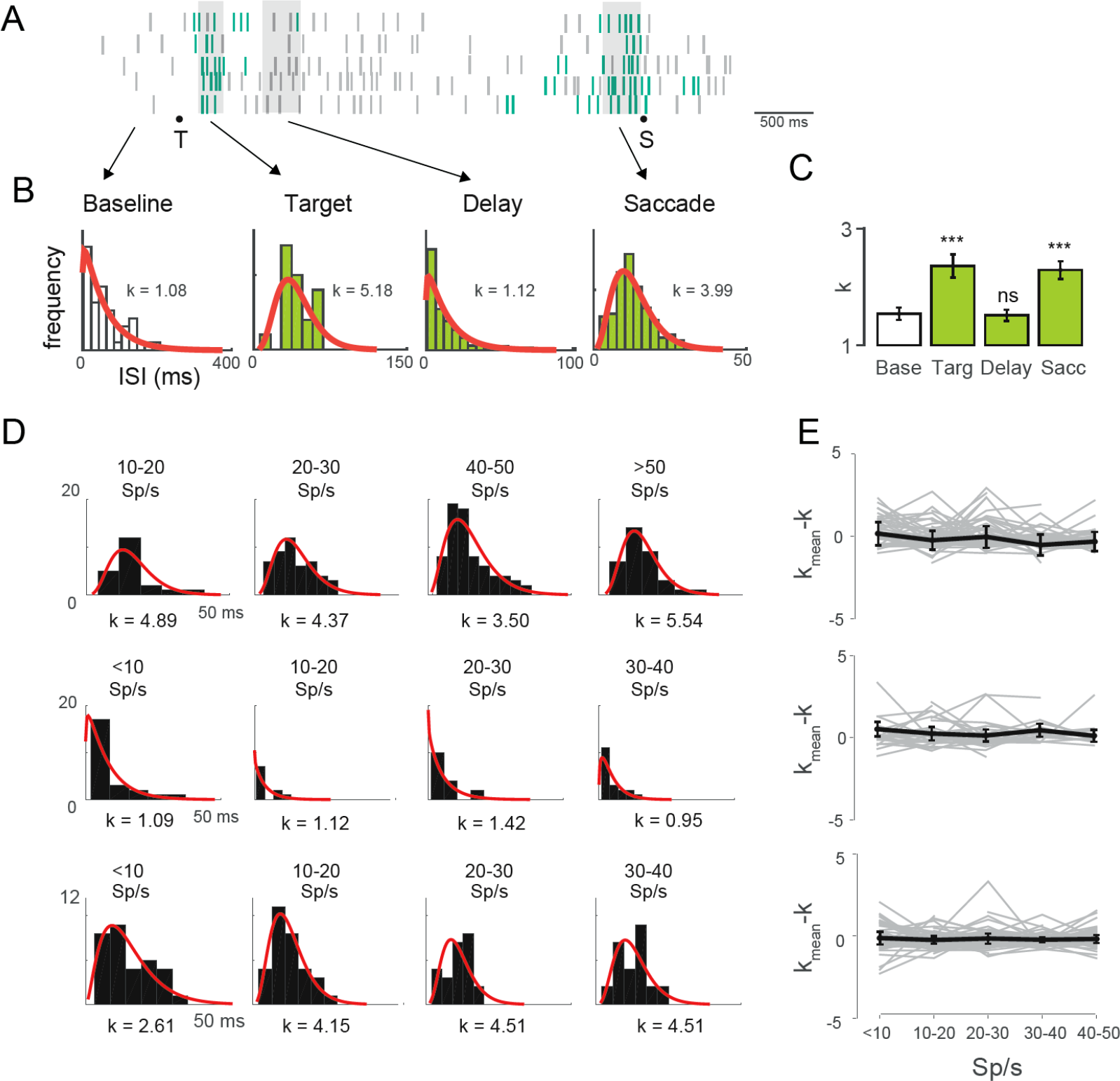
The decrease in within-trial variability in firing is independent of changes in firing rate. **A.** Raster plot of 5 trials of a representative neuron during target, delay and saccade epochs showing spikes that showed increased regularity and similar ISI (methods). **B.** The distribution of inter-spike intervals of spikes in the fixation (leftmost), target (left center), delay (right center) and saccade epochs (rightmost) fitted with a gamma distribution (red line). The shape parameter k, obtained from the fit is shown for the same representative neuron as 5A. **C.** The mean k values for all neurons for the four epochs. Statistics compares the k vales of rest of the three epochs with the fixation epoch value. *** means P<0.001 and n.s means not significant; ranksum test. **D.** Data for a representative neuron from A, separated based on bins of firing rate (as shown on the top of each panel) for the three epochs: target (top row), delay (middle row) and saccade epochs (bottom row). The k value for individual gamma fits (red line) is shown below each panel. **E.** Same analysis as in D, but k_mean – k values, for the population of neurons. None of the epochs showed significant modulation with firing rate (target epoch: P = 0.0871; delay epoch: P = 0.6251; saccade epoch: P = 0.8134, one way anova)

The firing rate increased in the target and saccade compared to the fixation and delay epochs. At higher firing rates, regularity in spiking could be merely due to a higher probability of occurrence of spikes. So, we next wanted to check if the effects we observed were merely due to this difference in firing rate. To test for spike regularity independent of underlying changes in firing rate, we computed rate-parsed histograms: For each cell, we plotted an ISI histogram for the interval associated with least firing rate in that epoch (typically 0-10 sp/s) and continued plotting ISI histograms for intervals of 10 sp/s up to the interval associated with the maximum firing rate of the cell in that epoch (typically 40-50 sp/s or rarely >50 sp/s). After generating a set of ISI histograms for each cell, for each of the three epochs (target, delay and saccade) we again fit these histograms to a gamma function and estimated their shape parameter, k. The ideal way to test if the spike regularity was independent of changes in firing rate would be to repeat the same analysis from Fig 4A-C within each interval of firing rates. However, this did not always give reliable results. This was because the firing rate in the baseline or delay epoch was at least three to four times lower than the firing rate in the target or saccade epoch and therefore, we rarely found any cell whose activity in any trial overlapped between the baseline/delay and the target/saccade epochs. Across the population, we found very few cells to show lower firing rates (<20 sp/s) during the target/saccade epochs and very few cells showing higher firing rates (>30 sp/s) during baseline/delay epochs. This potentially skewed the statistical test we performed due to higher degree of imbalance between the number of elements in the two groups that were tested. Nevertheless, we found (marginally) significant results (P values for target vs delay epoch: <10 sp/s: 0.0166; 10-20 sp/s: 0.0423; 20-30 sp/s: 0.0034; 30-40 sp/s: 0.0135; P values for saccade vs delay epoch: <10 sp/s: 0.0407; 10-20 sp/s: 0.0088; 20-30 sp/s: 0.0324; 30-40 sp/s: 0.0070; two-sample ttest or ranksum test). Additionally, to test this phenomenon in a different way, we performed an ANOVA within each epoch as a function of different rate intervals and we found no significant modulation in the shape parameter as a function of firing rate within a given epoch (target epoch: P = 0.0871, delay epoch: P = 0.6251, saccade epoch: P = 0.8134; one way ANOVA) indicating that the variation in spike rate did not confound our results (Fig 4 D-E).

Taken together, our results suggest that the spiking becomes more regular during neural events. In other words, the within-trial variability decreased during a stimulus driven response and movement preparation.

### Decrease in within-trial variability contributed to a decrease in across-trial variability

Both decrements in FF and increment in regularity of spiking indicate a deviation from Poisson firing. Here, we found evidence for both cases. Furthermore, the changes in the profile of average FF was highly correlated with that of CV2 (Fig 5, corr = 0.7912, P<10e-6). This already suggests that changes in FF could be due to changes in the underlying spike statistics ^24^. But do the changes in CV2 contribute to the changes in FF or are they just correlative? Since we couldn’t answer this directly through our dataset, we tested this hypothesis on simulated neurons. We simulated a stimulus neural event in 8 conditions (target positions), and presented them to 70 neurons, with different RF tunings, and drove them according to their tuning properties (see methods). We performed sets of simulations to test our hypothesis.

**Figure 5.**
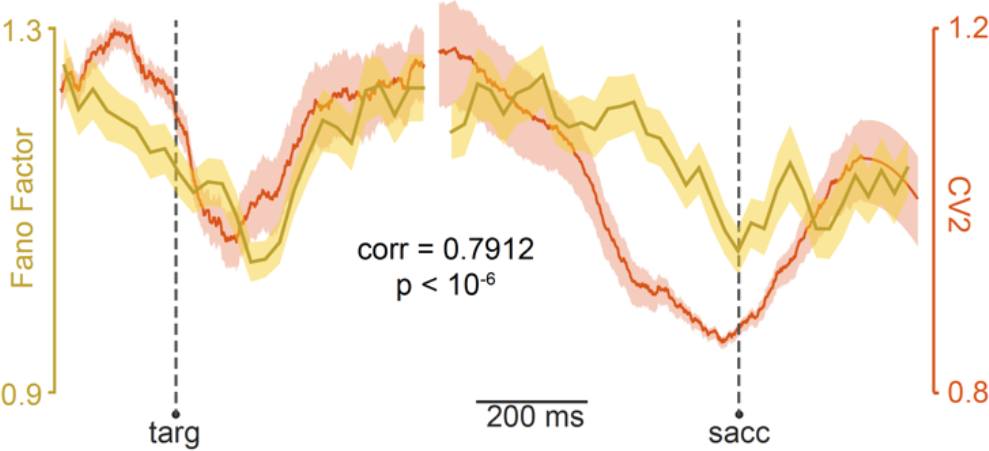
Changes in Fano Factor and CV2 are correlated across the population. Mean matched Fano factor (yellow) and average CV2 (red) are overlaid, aligned to target (left) and saccade (right). The profile of the two signals were highly correlated (corr = 0.7912, p<10e-6).

First, we increased the average firing rate of the population from a baseline value of ~10 sp/s to ~30 sp/s in response to the target onset, without changing the gamma shape parameter, keeping it at a constant at Poisson firing (k=1). This indicates a condition with irregular spiking or no change in within-trial variability, but with a change in firing rate in response to the target presentation. In this case, we did not see any change in the mean-matched FF (P = 0.6316; t-test). This shows that across-trial variability did not change (decrease) when there was no change in within-trial variability (Fig 6A). Note that, in terms of firing rate modulations, this scenario could either represent a case with no variability in true underlying rate or a constant variability in the underlying rate across trials. This is resolved in the next figure (Fig 7).

**Figure 6.**
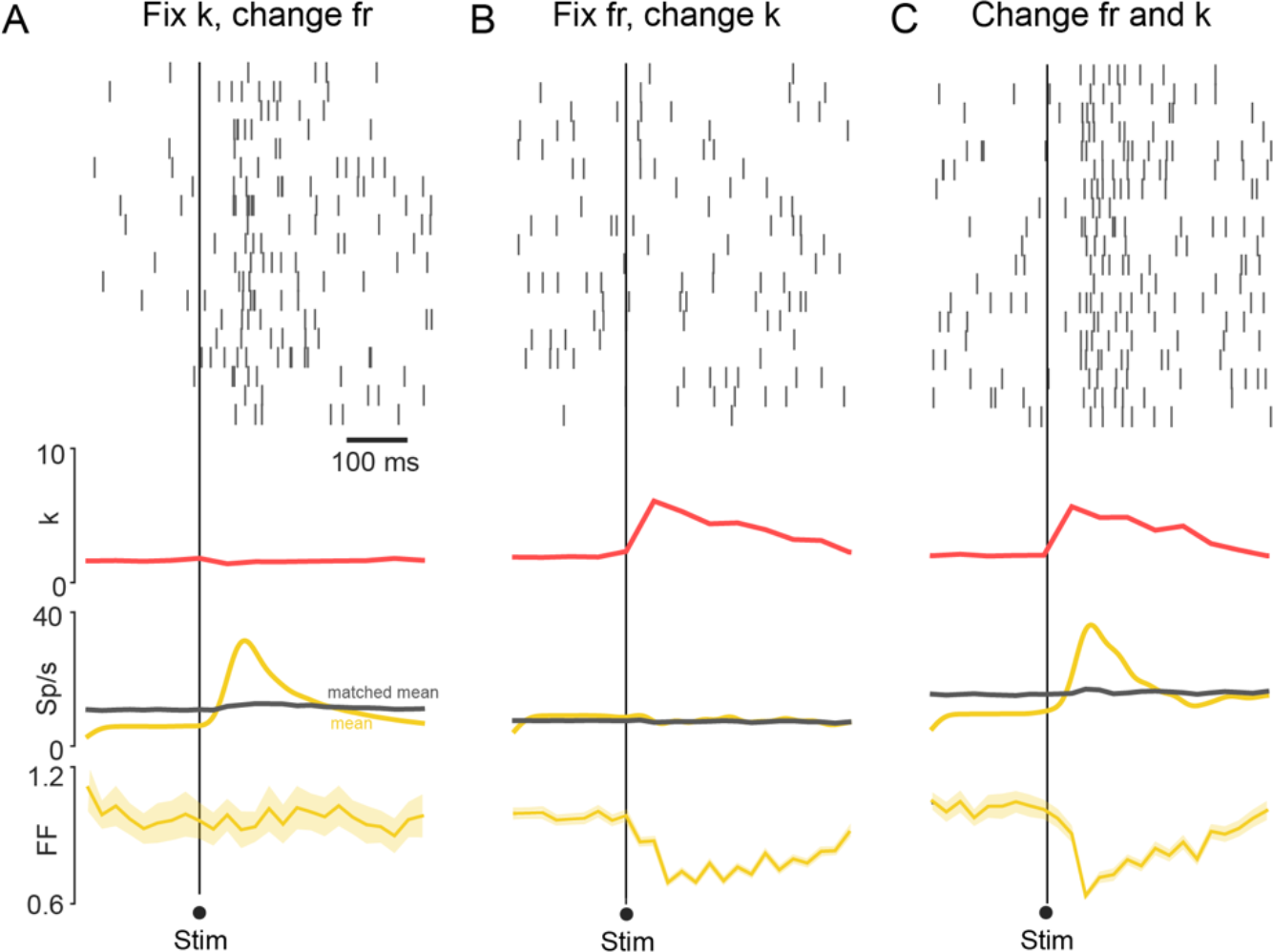
Contribution of within-trial variability to across-trial variability. **A:** Across-trial variability did not decrease when there was no change in within-trial variability: Top: Rasters aligned to stimulus onset for one condition, for a representative simulated neuron generated with no stimulus change in within-trial variability (k=1; Poisson model, red trace) but a stimulus dependent increase in firing rate (yellow trace). In this condition, there was no decrease in mean-matched FF (bottom panel). **B.** Across-trial variability decreased when there was a change only in the within-trial variability: Top: Rasters aligned to stimulus onset for one condition, for a representative simulated neuron generated with stimulus dependent reduction in within-trial variability (increase in k; red trace) but no stimulus dependent change in firing rate (yellow trace). In this condition, there was a decrease in mean-matched FF (bottom panel). **C.** Natural condition: Top: Rasters aligned to stimulus onset for one condition, for a representative simulated ‘neuron’ generated with stimulus dependent reduction in within-trial variability (increase in k; red trace) and a stimulus dependent increase in firing rate (yellow trace). In this condition, there was a further decrease in mean-matched FF (bottom panel).

**Figure 7.**
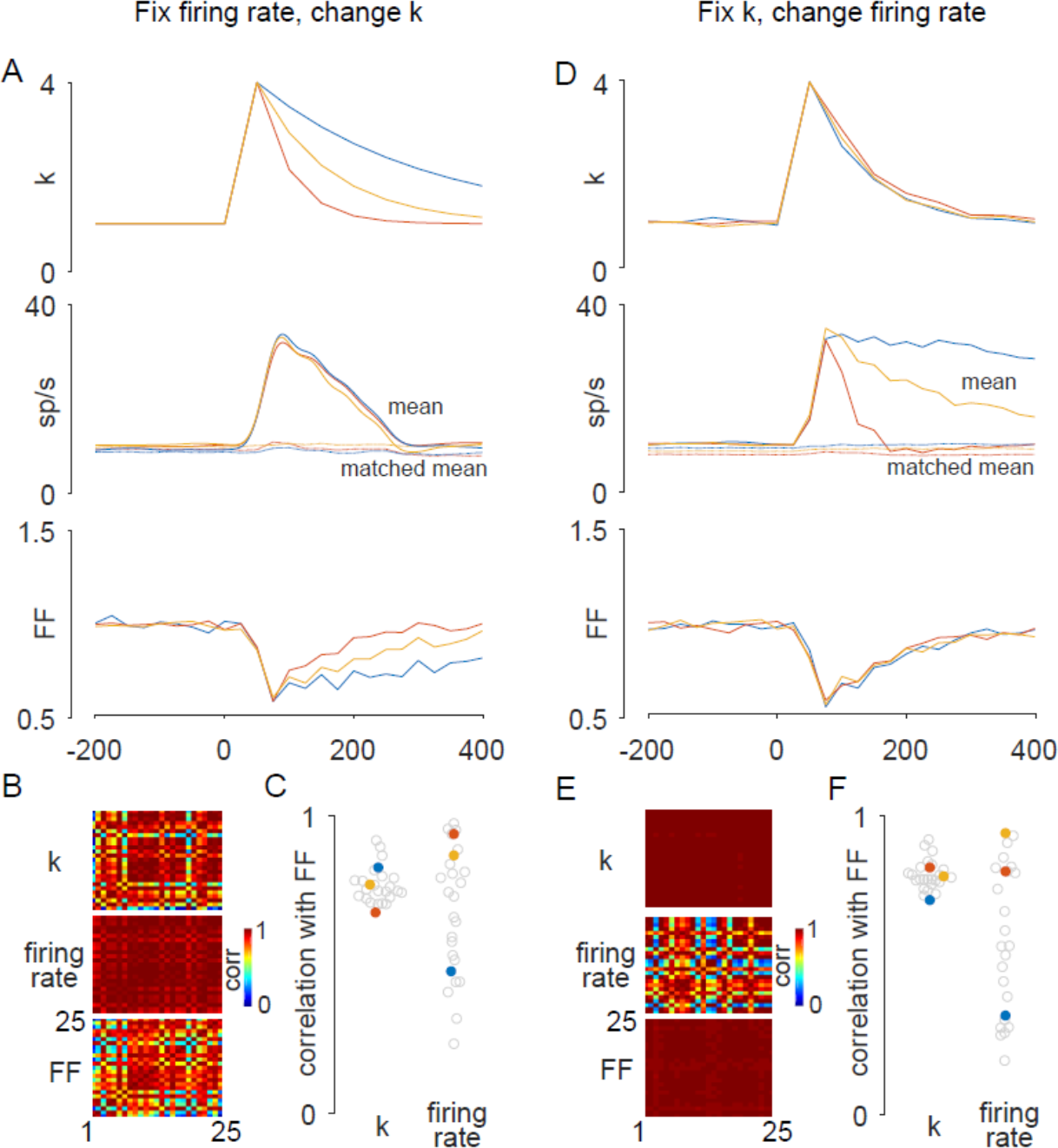
Time course of changes in within trial variability matches that of changes in across trial variability. **A**: Top: Three representative simulations with different time courses for the changes in within trial variability modelled per simulation as changes in the gamma shape factor, k. Middle: The firing rate corresponding to these three cases were modelled to be constant across simulations. Bottom: The calculated mean matched FF’s time course for these three cases. **B:** Time courses of each of the 25 simulations were correlated with themselves for k (top), firing rate (middle) and mean matched FF (bottom). **C**: Pearson correlation values of k (left) or firing rate (right) profile with the profile of Fano factor for each of the 25 simulations. The colored markers correspond to the sessions shown in A. **D:** Same as A but for constant within-trial variability profile across simulations and different firing rate profile per simulation. **E**: Same as B, but for simulations performed as mentioned in D. **F:** Same as C, but for simulations performed as mentioned in D.

Second, we only increased the gamma shape parameter, k from 1 (baseline Poisson) to ~3 after the target onset without changing the firing rate. This indicates a condition with regular spiking or a decrease in within-trial variability without a change in firing rate, in response to the presented target. As pointed out earlier, since at higher firing rates, regularity in spiking could be merely due to a higher probability of occurrence of spikes, we fixed the firing rate at a baseline level of 10 sp/s. In this condition, we observed a significant decrease in the mean-matched FF (P = 1.4750e-8; t-test) indicative of a decrease in the variability in true underlying rate. This shows that a decrease in within-trial variability could contribute to a decrease in across-trial variability (Fig 6B).

However, since we saw evidence for both increase in firing rate and regularity in spiking in our experimental results, we did a third simulation by increasing the firing rate (from ~10 sp/s to ~30 sp/s) and increasing k (from ~1 to ~3) in response to the target onset. Here, we saw a further reduction in mean-matched FF in this condition (P = 7.5345e-7; t-test; Fig 6C) again indicating a decrease in the variability in true underlying rate similar to our experimental findings.

Next, to further investigate the contribution of within trial variability to changes in across trial variability, we simulated 25 different conditions of 70 neurons each with different RF tunings. In each simulation, we simulated all the 70 neurons to have a simulation specific time course of change in within-trial spiking variability (by changing the shape factor k of the gamma distribution in every condition) in response to a presented target (Fig 7A **top panel**), while keeping their firing rate profile the same across all 25 simulations (Fig 7A **middle panel**). We then calculated the mean-matched FF for all 25 sessions independently (Fig 7A **bottom panel**) and found that the time course of the FF correlated well with the time course of the changes in within-trial variability and not the time course of changes in firing rate (Fig 7C). We correlated the entire simulated k and firing rate profiles and the calculated FF profile for each session with respect to other sessions to verify that the noise in the simulation process did not confound our result (Fig 7B; see methods). Then, we repeated this simulation but only changing the time course of the changes in firing rate every simulation, keeping the time course of changes in the within-trial variability the same across all simulations and then calculated the mean-matched FF for all 25 simulations independently (Fig 7D). Again, we found that only the time courses of the changes in within-trial variability correlated well with the time course of FF (Fig 7F). We did a similar correlation of the entire simulated k and firing rate profiles and the calculated FF profile for each simulation with respect to other simulation and verified that the noise in the simulation process did not confound our result (Fig 7E; see methods).

In conclusion, we reason that if the across-trial variability in neural firing is low, it suggests that the neuron fires with the same activity pattern across trials. But this could be possible if and only if the timing of spikes were approximately similar in that epoch (Fig 1B) on every trial. This cannot be achieved with a Poisson firing since the timing of firing is unpredictable (Fig 1A). Rather, if the firing were regular and stationary across trials, then there is a higher probability of spike timings being similar across trials thus reducing the variability across trials. Taken together, our results suggest that decrease in within-trial variability contributes significantly to a decrease in across-trial variability.

### Behavioral implication of decreased neural variability

A decrease in FF has been correlated with a faster reaction time (RT) in several cortical areas including motor cortices ^11^, lateral interparietal area ^7^, V4 ^14^ and frontal eye fields ^13^, among others. Our results suggest that this decrease in FF could have been due to increasing spike regularity. To test this possibility as a proof of principle, we calculated the FF and the CV2 for fast RT and slow RT conditions in the memory guided saccade task. Here, we calculated the RT as the time from go cue until the saccade onset (see methods). The mean RT across all sessions was 209.39 ± 0.31 ms (Fig 8A **top panel**) and did not vary as a function of the target location (P =0.999, one way ANOVA). For each session, the first quartile of RT was taken as ‘fast’ condition and the last quartile as the ‘slow’ condition. The fast RT and slow RT groups were significantly different across the population (fast RT: 122.75 ± 0.96 ms; slow RT: 311.99 ± 0.83 ms; P = 3.4761e-25, ranksum test; Fig 8A **bottom panel**). We found that both CV2 (P = 0.0073; ranksum test; Fig 8C) and mean matched FF (P = 0.0092; ranksum test; Fig 8B) in the saccade epoch were significantly lower for the fast RT condition compared to the slow RT condition.

**Figure 8.**
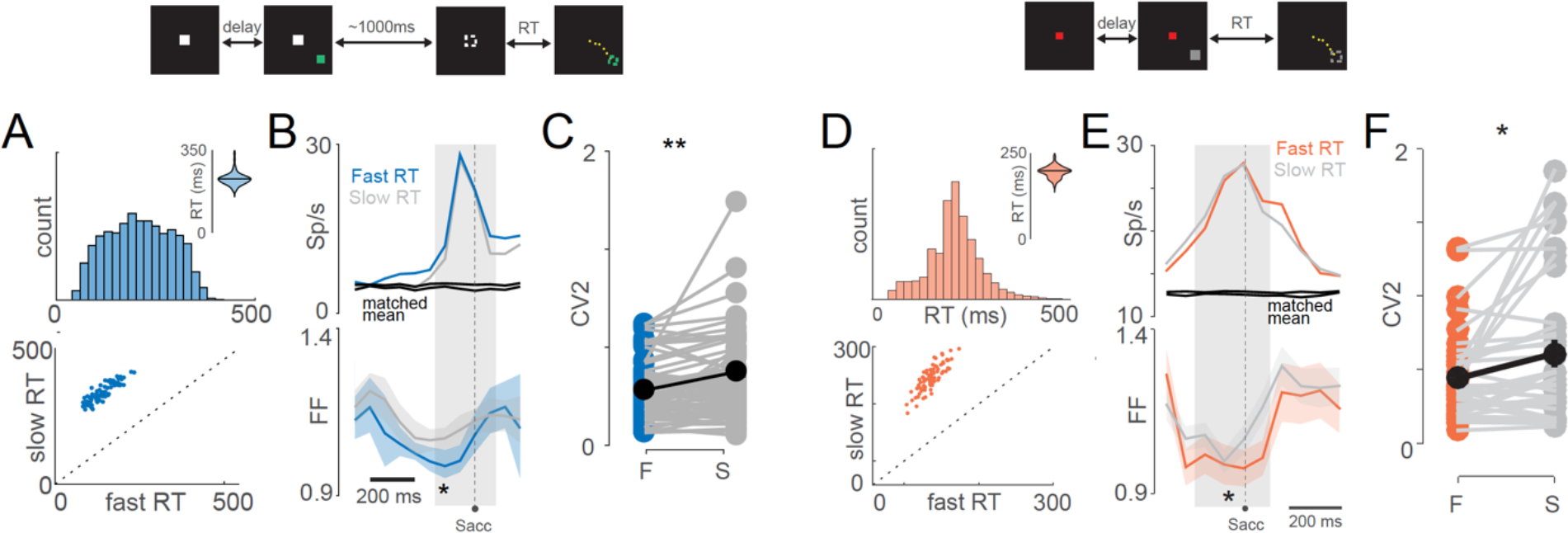
Reaction time correlates with within-trial variability in firing (proof of principle) **A:** Memory guided saccade task: Top: RT distribution across all sessions for all conditions Median = 210 ms. Inset: violin plot of median RT values of all sessions. Black horizontal line represents the mean: 209.39±0.31 ms. Bottom: median of fast RT vs slow RT plotted against each other for each session. **B**: Top: Mean spike density functions in the saccade epoch for fast RT (blue) and slow RT (grey) conditions. Mean-matched neural responses are shown in black. Bottom: Mean-matched FF for fast RT (blue) and slow RT (grey) conditions. * means P<0.05; ranksum test **C**: CV2 for fast RT (blue; F) and slow RT (grey; S) conditions for each neuron. ** means P<0.01; ranksum test **D:** Visually guided saccade task: Top: RT distribution across all sessions for all conditions Median = 200 ms. Inset: violin plot of median RT values of all sessions. Black horizontal line represents the mean: 196.10±0.40 ms. Bottom: median of fast RT vs slow RT plotted for each session. **E**: Top: Mean spike density function in the saccade epoch for fast RT (red) and slow RT (grey) conditions. Mean-matched neural responses are shown in black. Bottom: Mean-matched FF for fast RT (red) and slow RT (grey) conditions. * means P<0.05; ranksum test **F**: CV2 for fast RT (red; F) and slow RT (grey; S) conditions for each neuron. * means P<0.05; ranksum test

Additionally, we performed the same RT analysis in a visually guided saccade task also. Here, the monkeys had to make a saccade to a target flashed at one of the eight target locations as quickly as possible, still maintaining accuracy, to earn a liquid reward. The RT was the time elapsed between the target onset and the saccade initiation. The mean RT across all sessions was 196.10 ± 0.40 ms (Fig 8D **top panel**) and the mean RT did not vary as a function of the target location (P =0.999, one way ANOVA). The fast RT and slow RT groups were significantly different across the population (fast RT: 129.64 ± 0.37 ms; slow RT: 263.55 ± 0.59 ms; P = 1.7255e-47, paired t-test; Fig 8D **bottom panel**) We found that both CV2 (P = 0.0338; ranksum test; Fig 8F) and mean matched FF (P = 0.0139; ranksum test; Fig 8E) in the saccade epoch were significantly lower for the fast RT condition compared to the slow RT condition. Both these results provide a direct confirmation of our hypothesis that changes in within trial variability could contribute to changes in across trial variability.

## Discussion

Most cortical neurons either decode or encode information, or like visuomotor neurons of the FEF do both, as a part of the neural computations they instantiate along with other neurons. Although these computations are markedly different, the across-trial variability decreases in both these situations ^13^. In fact, this decline in across-trial variability is a common feature across different cortical areas ^4,11,13,14,17^ and changes in neural variability may be reflective of specific computations that occur in a brain area. For instance, changes in variability of FEF neurons is consistent with its active role in visual to saccade transformation ^18^. In this context, it is interesting to note that the decrease in variability during saccade planning is more prominent compared to the decrease in variability during the target information computation, suggesting that perhaps the latter computation is likely to be inherited from its inputs while saccade planning may be a more intrinsic computation of FEF ^13,18^. In this context, it is also interesting that the decrease in variability following target presentation is not maintained in FEF, but returns to baseline values, raising the possibility that signatures of variability may be more robustly expressed in more anterior regions of the PFC during the delay period ^7^. Similarly, changes in variability of PFC neurons, V4, LIP, premotor cortical neurons indicate cognitive engagement in a task ^12^, attention ^25^, salience to stimuli and reward ^7^, and motor preparation ^11^, respectively. Thus, measures of variability, in addition to measures of mean firing rates give valuable insights into a network level computation or population coding. However, since computations occur within and not across trials, it becomes important to have a principled approach to relate firing rate distributions to measures of firing rate activity within trials for measures such as FF to have meaningful value.

In this study, for the first time we have systematically related FF, the standard measure of across-trial variability, with CV2, a measure of within-trial variability, during behavioral epochs in which computations related to target encoding, working memory and motor preparation can be indexed within single neurons. We found a decrease in FF (Fig 2) after target presentation and during saccade planning epochs; and in the same epochs we found a decrease in CV2 (Fig 3) and an increase in the shape factor, k, of a gamma distribution fit to ISIs (Fig 4). Furthermore, the change in across-trial variability was highly correlated with the change in within-trial variability (Fig 5). We suggest that this decrease in across-trial variability in neural firing as indexed by FF, could be due to an increased spiking regularity or a decrease in within-trial variability in spiking, indexed by CV2 and k, during neural computations. We performed simulations of neurons to test whether there is a causal relationship between the two observations. We found that a decrease in within-trial variability leads to a decrease in across-trial variability during neural computations (Fig 6, 7). Finally, since a decrease in across-trial variability is linked to behavior such as reaction time, with decreased variability correlating with faster RT, we show that a faster RT is also correlated with an increased spiking regularity or a decrease in within-trial variability in spiking (Fig 8).

No previous study, according to our knowledge, has compared within and across-trial variability in such a systematic way. Furthermore, few computational studies have shown correlations between the two phenomena ^24,26^. Although it might seem self-evident that decreases in within-trial variability should decrease across-trial variability, this relation is not obvious, particularly since neurons are expected to inherit some of this variability due to shared network variability that may vary with time (Nawrot, Boucsein et al. 2008). We show a schematic of this aspect in Fig 1, where the within-trial variability could decrease without a decrease in the across-trial variability. However, since we empirically observed that across-trial variability in neural firing, was highly correlated with a decrease in within-trial variability in spiking, we conclude that spike train statistics are reasonably stationary across trials. One potential caveat in the current analyses is that due to the absence of simultaneous recording from multiple neurons, we are unable to determine whether the event based decrease in variability is due to changes in variability due to shared network variability or due to changes in spike generating mechanisms within individual neurons.

We found that although both FF and CV2 showed significant decrease from baseline variability both in and out of RF, there was a spatial modulation. Many studies show similar effects in the data although a discussion on the effect has been modest in the literature. Steinmetz and Moore showed that the response variability of V4 neurons changed as a function of the saccade direction ^14^. Falkner et al. showed that FF systematically decreases from the surround toward the saccade goal ^7^. In addition, they showed that LIP neurons displayed a dramatic decrease in FF only when the separation distance (between target and distractor) was >25°. Distances < 25° showed only a slight change in FF. In our dataset, the separation distance was 22° and this could potentially explain the observed spatial modulation. We did not vary the separation distance so we could not study the FF as a function of separation distance for FEF neurons. One other factor that could affect the spatial modulation of neural variability is the way in which the stimuli are presented. When Chang et al. presented only a single cue to a spatial location at a given time, FEF neurons showed a significant spatial modulation in FF between the RF and aRF conditions ^6^. But in the same experiment, when they presented a search array (six visual cues presented simultaneously in six equally spaced positions along an imaginary circle), they observed almost identical suppression in FF both in RF and aRF conditions. In relation to this, we only presented one cue at a time. Thus, this further explains the spatial modulation in neural variability in our data. Nevertheless, it should be noted that the underlying reasons for such effects on neural variability is currently unclear.

The observation that the presaccadic FF as well as CV2 are spatially tuned in FEF neurons implies only neurons that encode the endpoint of the upcoming saccade are reaching a minimum variance, which is in contrast to primary motor activity that exhibits a weak spatial tuning of response variability. While the lack of tuning of spatial variability in the motor cortex has been interpreted in support of an optimal subspace hypothesis in which all neurons in a cortical area initiate a movement when discharge rates converge to a specific value (Afshar et al. 2011), our data is consistent with previous work in FEF (Purcell, Heitz et al. 2012) in this regard. The observation that both within and across-trial variability are well tuned in FEF neurons implies that only neurons that encode the endpoint of the upcoming saccade are reaching a minimum variance and suggests that the optimal sub space hypothesis may not extend to oculomotor planning. Our results, however, are consistent with stochastic accumulator models of response preparation in which variability is expected to decline to a minimum at the time of the response. In this context, it is not surprising that like previous reports we also observed that the decrease in variability in both FF and CV2 prior to movement onset were correlated with changes in RT ^11,13,14^, indicating how decreases in variability influence behavior.

One way to reduce the neural variability in a network of neurons is to reduce the shared variance between neurons ^4,8^. Reductions in variability also enable neurons to be de-correlated and thus maximize the entropy or information content ^4^. Recent work looking at the activity of many neurons simultaneously have looked at the contributions on single trials have suggested a state space hypothesis in which variability shrinks at the start of the trial and this variability contributes to RT in a manner similar to FF and CV2. Interestingly this approach has also revealed novel features of activities (null space; oscillatory activity) that cannot be observed hitherto reported within single neurons ^4^. If within-trial variability is a major determinant of across-trial variability, as our work suggests, then a prediction from our work is that similar trajectories though neural space should be also discernable by random bootstrapping from the activity of single neurons across trials.

## Materials and methods

The dataset for this study comes from a previously published study ^18^. Please refer to that study for full details. We briefly describe the experimental procedures and methods here.

### Subjects

Two adult monkeys, J (male, *Macaca radiata*) and G (female, *Macaca mulatta*) were used for the experiments and were cared for in accordance with the Committee for the Purpose of Control and Supervision of Experiments of Animals (CPCSEA), Government of India, and the Institutional Animal Ethics Committee (IAEC) of the Indian Institute of Science. All protocols followed were in accordance with the guidelines set by the National Institutes of Health, USA.

### Data collection

TEMPO/VIDEOSYNC software (Reflective Computing, St. Louis, MO, USA) was used simultaneously with Cerebus data acquisition system (Blackrock Microsystems, Salt Lake City, UT, USA) for data collection. Eye positions were sampled with a mono-ocular infrared pupil tracker (ISCAN, Woburn, MA USA), interfaced with TEMPO software real time. The stimuli were presented on a calibrated Sony Bravia LCD monitor (21 inch; 60 Hz refresh rate) placed 57 cm in front of the monkey. Neural signals from 70 neurons were collected using single tungsten microelectrodes (FHC, Bowdoin, ME, USA; impedance: 2 to 4 MΩ) from the FEF on the right hemisphere through a permanently implanted recording chamber (Crist Instrument, Hagerstown, MD, USA).

### Task

#### Memory-guided saccade task

Details of this task have been described in detail elsewhere ^18^. The monkeys were required to fixate on a central fixation point (FP) for a variable amount of time at the start of the trial, following which a gray target appeared briefly at one of the eight equally-spaced peripheral locations on an imaginary circle of radius 12°. The monkeys had to continue fixation on the central FP for a variable delay (1,000 ms ± 15% jitter, sampled from a uniform distribution), following which the FP disappeared (go signal), cueing the monkeys to make a saccade to the remembered target location. On successful trials the monkeys received a juice reward.

For all further analyses, the target epoch was defined as VRL to VRL + 200ms after target onset where VRL is the visual response latency calculated by performing a t-test between the baseline (fixation epoch) and the signal after target onset ^18^. The delay epoch was defined as VRL + 500 ms to VRL+ 800 ms. The saccade epoch was defined as SPO to SPO+300 ms where SPO is the saccade plan onset which was calculated as the first point before the start of the saccade when the signal differed from baseline activity^18^. However, for visual FEF neurons or motor neurons where this calculation was not possible, the target epoch was taken as 50 ms to 200 ms after target onset, delay epoch was taken as 400 ms to 700 ms after target onset and the saccade epoch was taken as −200 ms to 100 ms centered on saccade onset.

#### Visually-guided saccade task

Each trial started with the appearance of a white central FP (0.6° × 0.6°). Following a variable time delay, a peripheral green target (1° × 1°) appeared in one of six possible locations on an imaginary circle of eccentricity 12°. The monkeys had to initiate a saccade as quickly and as accurately as possible to the target. On successful trials the monkeys received a juice reward.

Trials with artifacts including eye blinks, saccades with reaction times less than 100 ms on both tasks were removed prior to further data analysis.

#### Fano Factor analysis

The FF provides an estimate of the ‘noise to signal’ ratio of the data. We calculated FF by dividing the variance of spike counts by its mean in continuous bins throughout the trial (with a sliding window of 50 ms shifted by 10 ms). We calculated the mean-matched firing rate using the algorithm developed by ^4^.

#### CV2 analysis

CV2 compares adjacent ISI and calculates the level of similarity between them as a measure of intrinsic spiking variability. We calculated CV2 for i^th^ spike by computing the absolute difference in timing of two adjacent ISIs, divided it by their mean and multiplied it by 2 (see Fig 3A for an illustration).

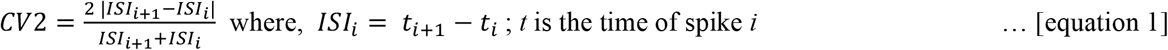

An average of CV2 over i spikes gives the variability of a given spike train, independent of the variations in firing rate in a short range (in the order of few milliseconds), since CV2 measures spiking variability in small intervals. Note that CV2 can only range from 0 (highly regular spiking; low within trial variability) to 2 (highly random spiking; high within trial variability) with a value of 1 indicating a Poisson firing (see Fig 3A for an illustration).

#### CV2 thresholding

To identify regular spiking patterns, we applied a threshold on the calculated CV2 values (Fig 4A). For a given set of spikes, if the CV2 value was less than or equal to the threshold, those ISIs were considered to be a part of a regular pattern with similar regularity in spiking (green spikes in Fig 4A). To do this, we measured the number of such patterns for a discrete range of CV2 values and set the threshold as the CV2 that resulted in maximum number of patterns ^27^. This analysis was just to illustrate that the spiking in target and saccade epochs has more regular patterned spiking than the fixation or delay epochs.

#### Gamma distribution fit

The gamma probability density function is a maximum entropy probability distribution and is defined by two parameters: shape (k) and scale (θ). Both these parameters are positive real numbers. The general form of the density function is given by the following equation:

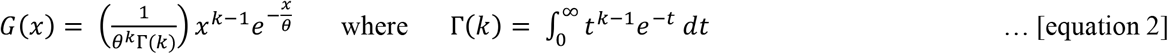

We calculated ISI histograms and fit them to gamma distributions (as shown above) using MATLAB’s gamfit function. This calculates the maximum likelihood estimates of k and θ After fitting the data, we performed a chi squared goodness of fit test that returned the decision for the null hypothesis that the data comes from a gamma distribution with parameters k and θ, estimated from the data. In specific cases where the data available per analysis was low, this test’s performance declined. Therefore, only in those cases, we visually inspected the fit for confirmation.^23^.

#### Simulations

We simulated 70 neurons with 8 conditions (target positions) per neuron and 20 trials per condition (to match our experimental dataset). For each neuron, four continuous conditions were randomly chosen to have a higher firing rate to the presented target compared to the other four conditions to mimic the RF-aRF property of neurons. We used the gamma probability density function (equation 2) with a shape parameter (*k*, that captures the shape of the inter-spike interval distribution) and scale parameter (θ, that approximates the firing rate) to generate spike trains with a firing rate, *r* given by:

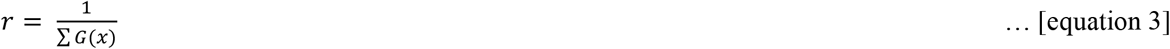

where *G*(*x*) is given by equation 2 above.

For each simulation, we fixed one of the two parameters: k, θ and we estimated the other parameter analytically or iteratively. Analytically,

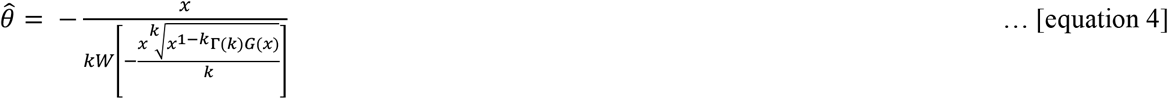

where W(z) is the product log function. However, there is no closed form solution to estimate k. Therefore, we first fixed a parameter (either k or θ) and fixed a value for *r*_*fix*_. Then, we iteratively sampled different values for the third parameter and calculated the firing rate of the spike trains thus generated from each of the gamma probability density functions (equation 2) and compared it to the firing rate that we fixed earlier, *r*_*fix*_. We repeated this process until the iteration converged to a solution. Finally, we used the mean-matching FF algorithm ^4^ to calculate FF on the matched mean.

#### Statistics

To check if two independent distributions were significantly different from each other, we first performed a two-sided goodness of fit Lilliefors test, to test for the normality, then used an appropriate t-test; or else a non-parametric Wilcoxon ranksum test. All values in this study, unless stated otherwise, are mean ± s.e.m.

## Author Contributions

NS: Designed the research, analyzed the data and wrote the paper.

DB: Collected the data, and wrote the paper.

AM: Designed the research and wrote the paper.

## Acknowledgements

This work was supported by an Intensification of Research in High Priority Areas Grant from the Department of Science and Technology, Government of India; a Department of Biotechnology - Indian Institute of Science (DBT-IISc) partnership programme grant; and institutional support from the Ministry of Human Resource Development.

